# Interactions among Merlin, Arkadia, and SKOR2 mediate NF2-associated Schwann cell proliferation in human

**DOI:** 10.1101/2024.09.24.614711

**Authors:** Pei-Ciao Tang, Seyoung Um, Anderson B. Mayfield, Olena R. Bracho, Christian Del Castillo, Christine T. Dinh, Derek M. Dykxhoorn, Xue Zhong Liu

**Author notes:** Equal contribution: Pei-Ciao Tang and Seyoung Um. Lead contact: Pei-Ciao Tang. Corresponding authors: Pei-Ciao Tang and Xue Zhong Liu.

## Abstract

NF2-Related Schwannomatosis (previously referred to as Neurofibromatosis Type 2, or NF2) is a genetic-associated disease resulting from mutations in the gene, *NF2*. *NF2* encodes the merlin protein, which acts as a tumor suppressor. Bilateral vestibular schwannoma (VS) is a hallmark of NF2. Although the exactly molecular mechanism mediating NF2-driven schwannomatosis remain unclear, it is known that defective Merlin protein functionality leads to abnormal cell proliferation. Herein, we utilized a human induced pluripotent stem cell (hiPSC)-based Schwann cell (SC) model to investigate the role of merlin in human SCs. SCs were derived from hiPSCs carrying a *NF2* mutation (c.191 T > C; p. L64P), its isogenic wild-type control cell line, and a NF2 patient-derived hiPSC line. NF2 mutant SCs showed abnormal cellular morphology and proliferation. Proteomic analyses identified novel interaction partners for Merlin – Arkadia and SKOR2. Our results established a new model in which merlin interacts with Arkadia and SKOR2 and this interaction is required for the proper activation of the SMAD-dependent pathway in TGFβ signaling.

## Introduction

Schwann cells (SCs) are the major type of glia cells and play crucial roles in the peripheral nervous system (PNS). SCs support the development, maintenance, and function of the PNS by myelinating axons and secreting trophic molecules(Jessen KR, 2005). SCs can be triggered by nerve injury to undergo cellular reprogramming and activate supportive functions, including the release of trophic factors and enhancing immune responses to promote neuron repair(Jessen and Mirsky, 2016). Abnormal SCs contribute to PNS disorders and injury. For example, schwannomatosis results from the formation of benign tumors, called schwannomas, on nerves. Although the factors that cause schwannomatosis are not fully understood, it is known that trauma in the PNS(Kennedy et al., 2016) and genetic mutations play important roles in schwannoma formation. Loss-of-function variants in the *NF2* gene (MIM 607379) are one of the most common drivers of schwannomas(Goetsch Weisman et al., 2023). Variants in *NF2* have been found in both sporadic and inherited forms of the disease(DG., 1998; Kluwe and Mautner, 1998). Furthermore, genetic variants in the *NF2* gene lead to a variety of nervous system tumors. Specifically, bilateral vestibular schwannoma (VS), benign tumors resulting from the neoplastic growth of SCs of the vestibulocochlear nerves, is a major diagnostic criteria for *NF2*-related schwannomatosis (previously referred to as Neurofibromatosis type 2, or NF2)(Dinh et al., 2020; Plotkin et al., 2022). Although benign, VSs can involve the vestibulocochlear nerves and cause hearing loss and balance problems. The *NF2* gene encodes the tumor surpressor merlin (Moesin-Ezrin-Radixin-Like Tumor Suppressor) protein that is involved in many signaling pathways depending on the specific tumor types, including the Hippo signaling pathway, WNT/β-catenin signaling pathway, TGFβ signaling pathway, and receptor tyrosine kinase signaling to serve as a tumor suppressor(Goetsch Weisman et al., 2023; Mota and Shevde, 2020; Nourbakhsh and Dinh, 2023). For example, merlin phosphorylation at p.S518 by the p-21-activated kinase 2 (PAK2) is regulated by TGFβ signaling in epithelia(Kissil et al., 2002; Wilkes et al., 2009) and the capability for Merlin binding to PAK (PAK1 and 2) alters merlin tumor suppressor function(Kissil et al., 2003; Wilkes et al., 2009; Xiao et al., 2005).

Although variants in *NF2* are the major genetic drivers of the formation of schwannomas, the molecular mechanisms by which *NF2* mutations drive abnormal SC proliferation are still not fully understood. Previous studies on the function of Merlin have provided invaluable insights into its cellular roles. However, many of these studies were carried out in non-human SC systems, including mouse models and human immortalized cell lines(Curto and McClatchey, 2008),(Chalak M, 2024; McClatchey AI, 1997). Stem cell-based models provide an alternative to transformed cells since they maintain the genetic architecture of the human genome and the genetic susceptibility to disease. Here in, we have used the human induced pluripotent stem cells (iPSC)-derived SC system(Kim HS, 2017; Majd H, 2023) to model the formation of schwannomas and the molecular mechanisms that govern this process. Specifically, we showed that hiPSC-derived SCs bearing patient-specific variants in NF2 recapitulate the abnormal cell proliferation phenotype seen in schwannomas(Gutmann DH, 1998).

Furthermore, proteomic analyses were performed to investigate the role of merlin in cell proliferation. We identified novel merlin interaction partners, Arkadia and SKOR2, and show that the L64P (c.191 T>C; p.L64P) variant disrupted these protein-protein interactions. Disrupting these interactions altered the response to the TGFβ signaling pathway. The patient deletion bearing iPSC-derived SCs validated the role of this mechanism in driving cellular proliferation. Through these approaches, we elucidated the molecular mechanisms underlying the abnormal proliferation resulting from the *NF2* mutations in SCs and proposed a novel mechanism by which merlin suppresses SC overgrowth.

## Results

### Differentiation of NF2^WT^ and NF2^L64P^ hiPSC lines into SCs

We previously established a hiPSC line carrying a homozygous patient-specific *NF2* mutation, p.L64P(Nourbakhsh et al., 2021). SCs were differentiated from both the NF2^L64P^ and its isogenic wildtype control parental cell line NF2^WT^ using a previous published protocol(Kim HS, 2017) with modifications (Figure 1A). Both NF2^WT^ and NF2^L64P^ underwent the first induction phase and generated Schwann cell progenitor (SCPs) that had comparable cell numbers as assessed by the expression of SOX10 (Figure 1B-B”) by the total induction day 18 without any noticeable differences in cell morphology. By day 14 of the maturation stage (the total induction day 32), we observed the induction of SCs based on S100β signals with approximately 80% of total cells staining S100β+ in both NF2^WT^ and NF2^L64P^ cell lines (Figure 1C-C”). In addition to SCs, these cultures contained ~5-10% neurons based on the NeuN staining (Figure 1D-D”).

**Figure 1.**
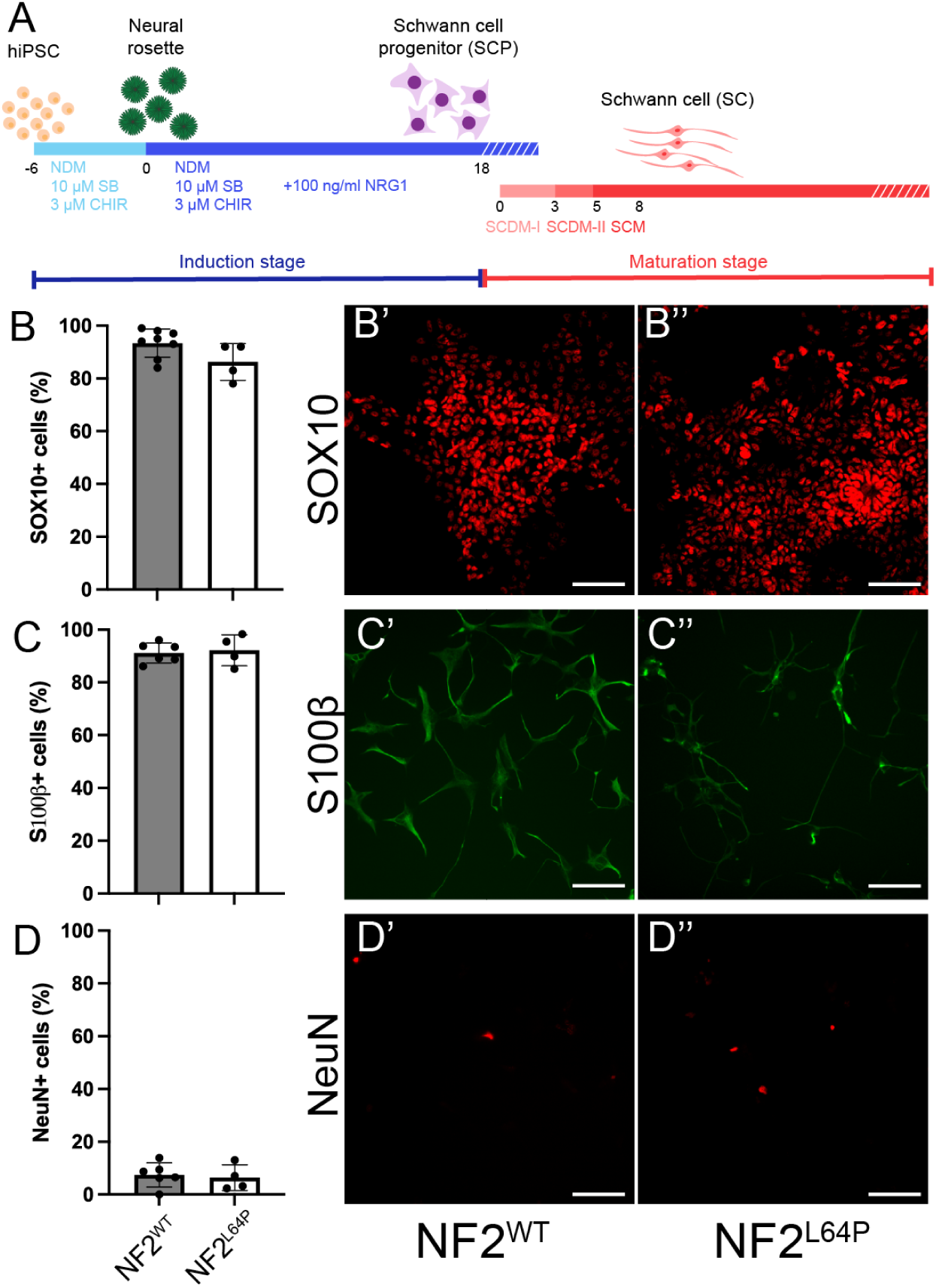
Differentiation of Schwann cells (SCs) from NF2^WT^ and NF2^L64P^ hiPSC lines. A. Schematic of the differentiation protocol. B, C, and D. Quantification of Schwann cell progenitors (B), SCs (C) and neurons (D) induction based on the ratio of marker+ cells over DAPI+ cells. B’, B”, C’, C”, D’ and D”. Representative immunohistochemistry (IHC) images of markers for SCPs, SCs, and neurons, respectively. Scale bar = 100 µm.

### Higher cell proliferation level and abnormal cell morphology in NF2^L64P^-derived SCs

Interestingly, proliferation was significantly higher in the NF2^L64P^ hiPSC-derived SCs compared to NF2^WT^ hiPSC-derived SCs based on BrdU incorporation assays on the day 14 of the maturation stage (Figure 2A). Additionally, we observed distinct morphological differences between the NF2^L64P^ SCs and the NF2^WT^ SCs (Figure 2B-B’). Specifically, SCs derived from the NF2^WT^ hiPSC line exhibited a bipolar shape with small ruffles at the ends of two poles (Figure 2B). On the other hand, NF2^L64P^ hiPSC-derived SCs lacked this polarity and, instead, exhibited a more “spreadout” cell shape with larger ruffles and a greater cytoplasmic volume (Figure 2B’). Indeed, measurements of cell area indicated significantly larger cell size in NF2^L64P^ hiPSC-derived SCs than that seen in NF2^WT^ hiPSC-derived SCs (Figure 2C-C”). Our findings demonstrated that the NF2 mutation, p.L64P, bearing hiPSCs-derived SC results in phenotypes consistent with observation reported in previous studies, including elevated SC proliferation and alterations in the cell morphology(Gutmann DH, 2001; Gutmann et al., 1999).

**Figure 2.**
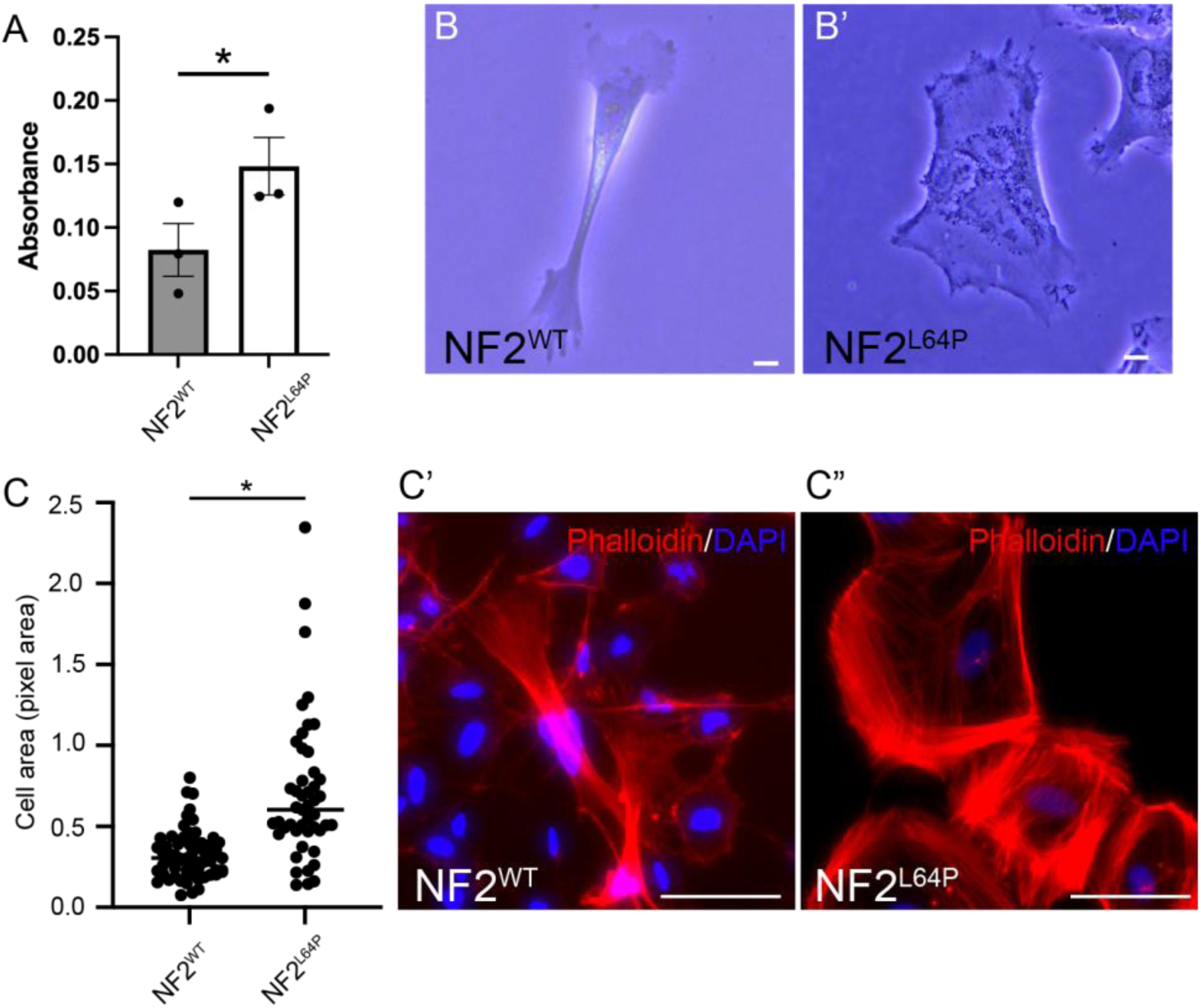
Abnormal phenotypes in NF2^L64P^-derived Schwann cells (SCs). A. Higher Cell proliferation level in NF2^L64P^-derived SCs based on the BrdU assay. B and B”. Representative bright-field images of SCs derived from NF2^WT^ and NF2^L64P^ hiPSC lines. Scale bar = 10 µm. C. Significantly larger cell surface area in NF2^L64P^-derived SCs. C’ and C”. Representative IHC images of SCs derived from NF2^WT^ and NF2^L64P^ hiPSC lines staining with phalloidin for F-actin. Scale bar = 100 µm.

### The *NF2* mutation, p. L64P, significantly alters the proteome of hiPSCs-derived SCs

In order to investigate the role of merlin in SC biology, we performed co-immunoprecipitation (Co-IP) analysis. The merlin protein was precipitated from protein lysates isolated from NF2^WT^ and NF2^L64P^ hiPSC-derived SCs on the maturation day 14 using a merlin-specific antibody (Figure 3A). The resulting merlin-associated proteins were analyzed by SDS-PAGE gel electrophoresis followed by imaging using the Bio-Rad Stain free gel imaging system. Interestingly, there were distinct patterns of protein banding observed in the NF2^L64P^ compared to the NF2^WT^ samples suggesting that the L64P mutation alters the binding properties of merlin (Figure 3A). Tandem mass tags (TMT), coupled with liquid chromatography–tandem mass spectrometry (LC–MS/MS) proteomic analysis was performed to identify the differential sets of proteins bound to the WT and L64P variant bearing versions of merlin.

**Figure 3.**
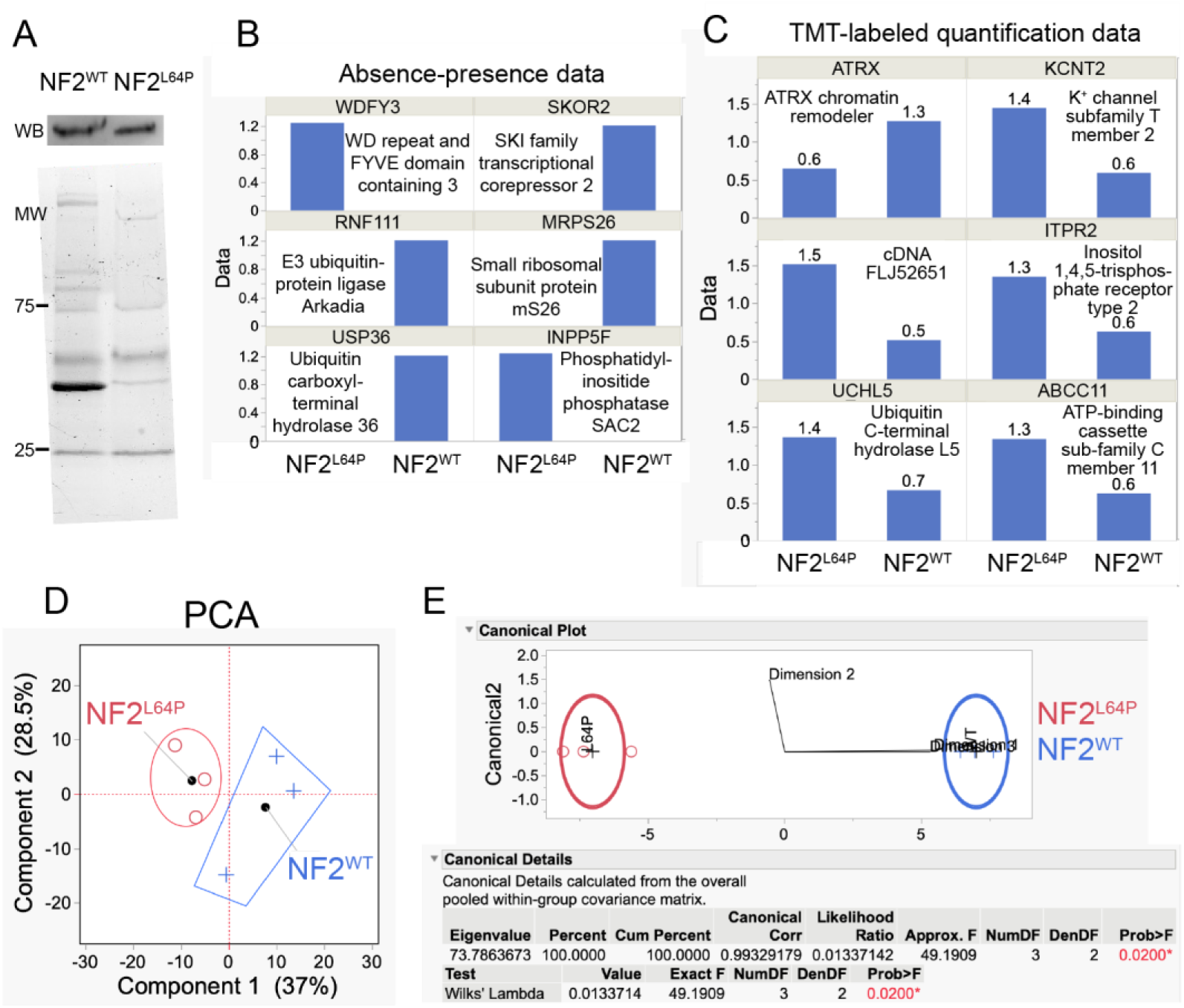
Proteomic analyses of proteins after co-immunoprecipitation (Co-IP) with the Merlin antibody from NF2^WT^- and NF2^L64P^-derived SCs. A. Different protein pattern between NF2^WT^ and NF2^L64P^ after IP in SDS-PAGE with the Merlin antibody that recognizes both WT and mutant Merlin. B. Six differentially concentrated proteins (DCPs) were identified via analyzing presence-absence data. C. Six DCPs were identified in the TMT-based quantification data. D. TMT-based data showed distinctions in proteomes between samples from NF2^WT^ and NF2^L64P^ in the principal component analysis (PCA). E. Canonical analysis suggested strong effects of the NF2 p.L64P mutation on SC proteome.

Using the presence-absence approach after data normalization (Figure S1A), six differentially concentrated proteins (DCPs) were identified (Figure 3B) out of a total of 1,076 proteins (of which 621 [58%] were housekeeping proteins) identified. Of these six (~1% of the 455 proteins whose levels varied across the six samples), four were only found in the NF2^WT^ samples (Figure 3B) while the other 2 proteins were only found bound to the L64P variant bearing merlin. The NF2^WT^-associated proteins included a SKI family transcriptional corepressor 2 (Uniport ID: Q2VWA4), an E3 ubiquitin protein ligase (Uniport ID: Q6ZNA4), a small ribosomal subunit protein mS26 (Uniport ID: Q9BYN8), and a ubiquitin carboxyl-terminal hydrolase (Uniport ID: Q9P275). The proteins bound exclusively to the NF2^L64P^ protein were the WD repeat and FYVE domain containing 3 protein (Uniport ID: A0A1D5RMR8) involved in autophagy, and the phosphatidylinositide phosphatase SAC2 (Uniport ID: Q9Y2H2). None of the associated peptides were labeled with TMT. Though a discrimination analysis was not significantly different, samples from NF2^WT^ and NF2^L64P^ nevertheless were well separated in the canonical plot base on their protein profiles (Figure S1B).

When looking only at a subset of 262 peptides that were labeled with TMT after data normalization (Figure S1C), an additional six DCPs were identified (Figure 3C). Of these, only an ATRX chromatin remodeler (Uniport ID: A0A096LNX6) was maintained at higher levels in the NF2^WT^ (2.2-fold). The remaining five were found at high levels only in the NF2^L64P^-associated proteomes: a potassium channel subfamily T member 2 (Uniport ID: A0A6E1ZGS3; 2.3-fold), an unknown protein encoded by cDNA FLJ52651 (Uniport ID: B7Z8Y8; 3-fold), an inositol 1,4,5-trisphosphate receptor type 2 (Uniport ID: F5GYT5; 2.2-fold), a ubiquitin C-terminal hydrolase L5 (Uniport ID: Q5LJB1; 2-fold), and an ATP-binding cassette sub-family C member 11 (Uniport ID: Q96J66; 2.2-fold).

Principal component analysis (PCA) biplot explained ~2/3 of the variation in the TMT dataset across the first two PCs and some clustering by treatment is evident in Figure 3D. To quantify this difference, a discriminant analysis of the first three multidimensional scaling (MDS) coordinates was undertaken (NP-MANOVA; i.e., discriminant analysis of genotypes), and a statistically significant Wilks’ lambda was obtained (*p*=0.02; Figure 3E). This means that the partial Co-IP proteomes of the NF2^WT^ and NF2^L64P^ protein differed significantly from one another, although only 6 of 262 TMT-labeled proteins (~2%) were deemed DCPs by our conservative, dual-criteria approach.

### Merlin interacts with Arkadia and SKOR2 and such interaction mediates the degradation of SKOR2 in nuclei

The E3 ubiquitin ligase Arkadia has been previously shown to ubiquitinate members of the SKI family of proteins leading to their degradation by the ubiquitin–proteasome system (UPS) resulting in enhanced TGF-β signaling(Briones-Orta et al., 2013; Levy et al., 2007). Since both Arkadia and the SKI family member SKOR2 showed differentially binding between NF2^WT^ and NF2^L64P^ proteins, we examined whether Arkadia could regulate SKOR2 function in a merlin-dependent manner. We hypothesized that merlin interacts with Arkadia to induce SKOR2 degradation. To begin, we validated the binding between merlin and Arkadia and SKOR2 via Co-IP followed by Western blot (Figure 4A and B). This interaction was significantly diminished by the L64P mutation in *NF2* (Figure 4A). Further, we demonstrated that immunoprecipitation using an antibody against Arkadia led to the pull down of SKOR2 and merlin (NF2) (Figure 4B) and results confirmed the interactions between merlin, Arkadia, and SKOR2. Intriguingly, we noticed that, instead of a band with the predicted size at approximately 105 kDa as was observed in the whole lysate samples (Figure S2), several bands at smaller sizes were detected in the Western blots against SKOR2 after the Co-IP (Figure 4A-B). As previously mentioned, Arkadia was reported to ubiquitinate SKI family proteins for the subsequent protein degradation, we also performed the Western blot with the ubiquitin antibody following the Co-IP with merlin. Interestingly, a similar band pattern to the Western blots of SKOR2 after Co-IP was seen (Figure 4C). These results suggest that SKOR2 protein interacting with merlin and Arkadia was likely being degraded through the UPS.

**Figure 4.**
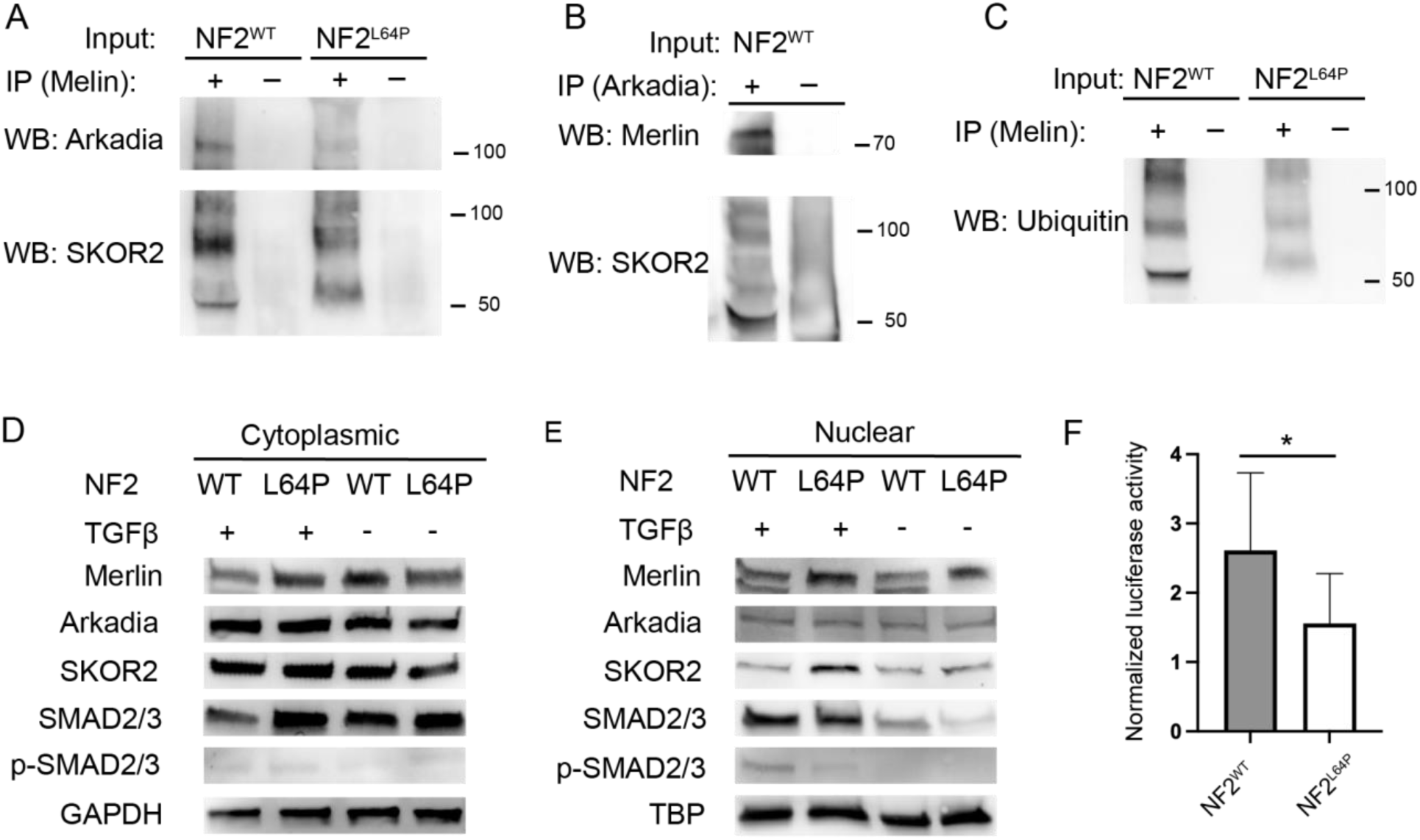
Merlin interacts with Arkadia and SKOR2 in the SMAD-dependent pathways in the TGFβ signaling. A. Western blots of Arkadia and SKOR2 following the Co-IP with the merlin antibody. B. Western blots of Merlin and SKOR2 following the Co-IP with the Arkadia antibody. C. Similar Western blot pattern of Ubiquitin following the Co-IP with the merlin antibody with the Western blots of SKOR2 in panel A and B. D. Western blots in the cytoplasmic fraction. E. Western blots in the nuclear fraction. F. The SBE assay indicated significantly higher response to the TGFβ activation in NF2^WT^-SCs comparing to its in NF2^L64P^-derived SCs.

After confirming that merlin binds to Arkadia and SKOR2, we then investigated whether this interaction affects the degradation of SKOR2 to regulate TGFβ signaling. We evaluated the presences of key proteins in the TGFβ pathway – merlin, Arkadia, SKOR2, and phosphorylated SMAD 2 and 3 (p-SMAD2/3) – in cytoplasmic and nuclear protein fractions from both NF2^WT^- and NF2^L64P^ hiPSC-derived SCs. There was no obvious difference in the level of these proteins in whole lysates (Figure S2) and cytoplasmic protein fractions (Figure 4D) isolated from NF2^WT^ and NF2^L64P^ SC. However, there was significantly lower levels of SKOR2 in the nuclear protein fraction isolated from the NF2^WT^ hiPSC-derived SCs compared to that from the NF2^L64P^ hiPSC-derived SCs (Figure 4E). Moreover, there was stronger p-SMAD2/3 levels only in SCs-derived from NF2^WT^ hiPSCs treated with TGFβ (Figure 4E), though there were no differences in SMAD2/3 signal in the cytoplasmic fraction (Figure 4D) and the equivalent level of translocation of SMAD2/3 in the nuclear fraction was seen between NF2^WT^ and NF2^L64P^ samples with TGFβ activation (Figure 4E). To functional test the activity of TGFβ/SMAD signaling pathway in the NF2^WT^ and NF2^L64P^ iPSC-derived SCs, the SBE assay was performed. The SBE reporter assay is a SMAD-dependent TGFβ pathway-responsive luciferase reporter assay. We found significantly higher SBE activity in the NF2^WT^ compared to the NF2^L64P^ hiPSC-derived SCs.

Overall, our data suggested that wild-type merlin protein is required for the degradation of SKOR2 in the nuclei and, further, the stability of SKOR2 is critical for the SMAD-dependent response to TGFβ activation. The p.L64P mutation in the merlin protein disrupted this function and, ultimately, altered the TGFβ signaling pathway.

### Patient-specific iPSC-derived SC model validates the role of merlin in cell proliferation and TGFβ signaling

An hiPSC line was derived from peripheral blood mononuclear cells (PBMCs) isolated from a patient bearing a heterozygous deletion in chromosome 22, including the *NF2* gene (NF2^+/−^) (Figure S3). Global screening array (GSA) confirmed the partial deletion in chromosome 22 in the hiPSC line (Data not shown). SOX10+ SCPs were generated from the NF2^+/−^ hiPSC line (Figure 5A-A’) and subsequently differentiated into S100β+ SCs (Figure 5B-B’). Phalloidin staining showed that the polarized F-actin distribution in the NF2^+/−^ and NF2^WT^ SCs (Figure 5C-C’). Although equivalent numbers of cells were plated and the same culture conditions were used, we consistently observed more cells in the NF2^+/−^ SC compared to that of the NF2^WT^ cultures (Figure 5C-C’). To determine if this discrepancy in cell number was due to elevated levels of cell proliferation, BrdU incorporation was measured in the NF2^+/−^ and NF2^WT^ on the total induction day 32 (day 14 of the maturation stage) cultures. Indeed, the NF2^+/−^ hiPSC-derived SCs exhibited significantly higher cell proliferation activity than the NF2^WT^ hiPSC-derived SCs (Figure 5D).

**Figure 5.**
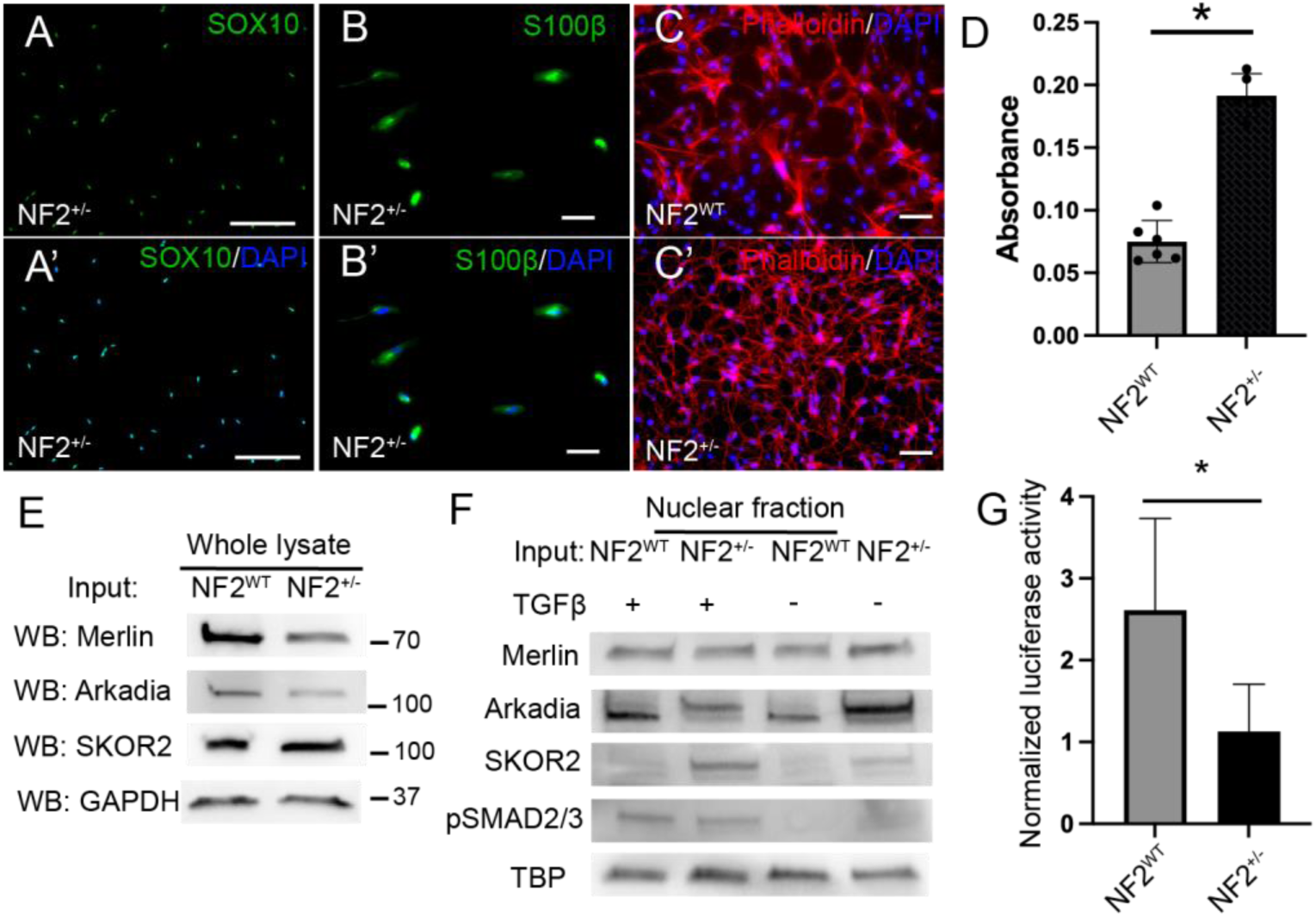
Abnormal cell proliferation in SCs derived from the patient-derived hiPSCs (NF2^+/−^). Similar Western blot results among merlin, Arkadia, SKOR2, pSMAD2/3 in nuclear fractions in responding to the TGFβ activation indicated that heterozygous loss of *NF2* results in the higher SKOR2 and the significantly lower SBE activity level. A-A’. Representative IHC images of SOX10+ SCPs derived from NF2^+/−^ hiPSCs. B-B’ Representative IHC images of S100β+ SCs derived from NF2^+/−^ hiPSCs. C-C’ Representative images of NF2^WT^ hiPSCs- and NF2^+/−^hiPCSs-derived SCs staining with phalloidin. D. Significantly higher proliferation activity in NF2^+/−^-derived SCs. E. Western blots in whole lysates. F. Higher level of SKOR2 in NF2^+/−^-derived SCs with the TGFβ activation. G. Significantly lower SBE activity in NF2^+/−^-derived SCs comparing NF2^WT^-derived SCs after the TGFβ activation.

To determine if the heterozygous NF2 deletion altered TGFβ signaling in a manner similar to that seen with NF2^L64P^ hiPSC-derived SCs, we analyzed the level of merlin, Arkardia, SKOR2, p-SMAD2/3, and SBE activity following TGFβ activation. Similar to the results seen with the NF2^L64P^ hiPSCs-derived SCs, there were no discernible difference in protein levels in the whole cell lysates isolated from NF2^WT^- and NF2^+/−^ hiPSCs-derived SCs (Figure 5E). There was a noticeable lower signal for merlin in the NF2^+/−^ sample compared to the NF2^WT^ samples (Figure 5E) as would be expected since the NF2^+/−^ sample lacks one copy of this gene. Similar to the results seen in the NF2^L64P^ SCs, the heterozygous NF2 deletion SCs (NF2^+/−^) had higher levels of SKOR2 in the nuclear protein fraction compared to the NF2^WT^ SCs (Figure 5F). In addition, the NF2^+/−^ SCs had decreased p-SMAD2/3 levels compared to the NF2^WT^ SCs (Figure 5F) suggesting a decrease in TGFβ signaling. To confirm that the NF2^+/−^ SCs had decreased TGFβ signaling, SBE activity was measured in NF2^+/−^ hiPSC-derived SCs compared to NF2^WT^ SCs. Consistently, the NF2^+/−^ SCs had significantly lower SBE activity compared to the NF2^WT^ SCs (Figure 5G). These results support our findings and hypothesis that NF2 acts as a modulator of TGFβ signaling through its interaction with Arkadia and SKOR2 as was seen in the NF2^L64P^ SCs. Furthermore, although the NF2^+/−^ SCs didn’t show the same alteration in cellular morphology as the NF2^L64P^ cells, NF2^+/−^ still showed a deficit in merlin, Arkadia, and SKOR2 are interaction and TGFβ signaling suggesting that merlin contributes to the maintenance of adequate cell proliferation in human SCs.

## Conclusion

In this study, our results suggest a model in which merlin is required for Arkadia to ubiquitinate SKOR2 (Figure 6). The p. L64P mutation was shown to disrupt this interaction, allowing SKOR2 to accumulate in the nucleus (Figure 6). Degradation of SKOR2 is necessary for the activation of TGFβ-responsive gene expression by phosphorylated SMAD proteins, i.e., p-SMAD2/3 (Figure 6), which may be important for regulating cell proliferation. These findings were further supported by the results from the NF2 patient-derived hiPSCs carrying a heterozygous deletion of the *NF2* gene. In conclusion, we propose a new model of merlin activity as a tumor suppressor through our identification of novel protein-protein interaction partners in human SCs.

**Figure 6.**
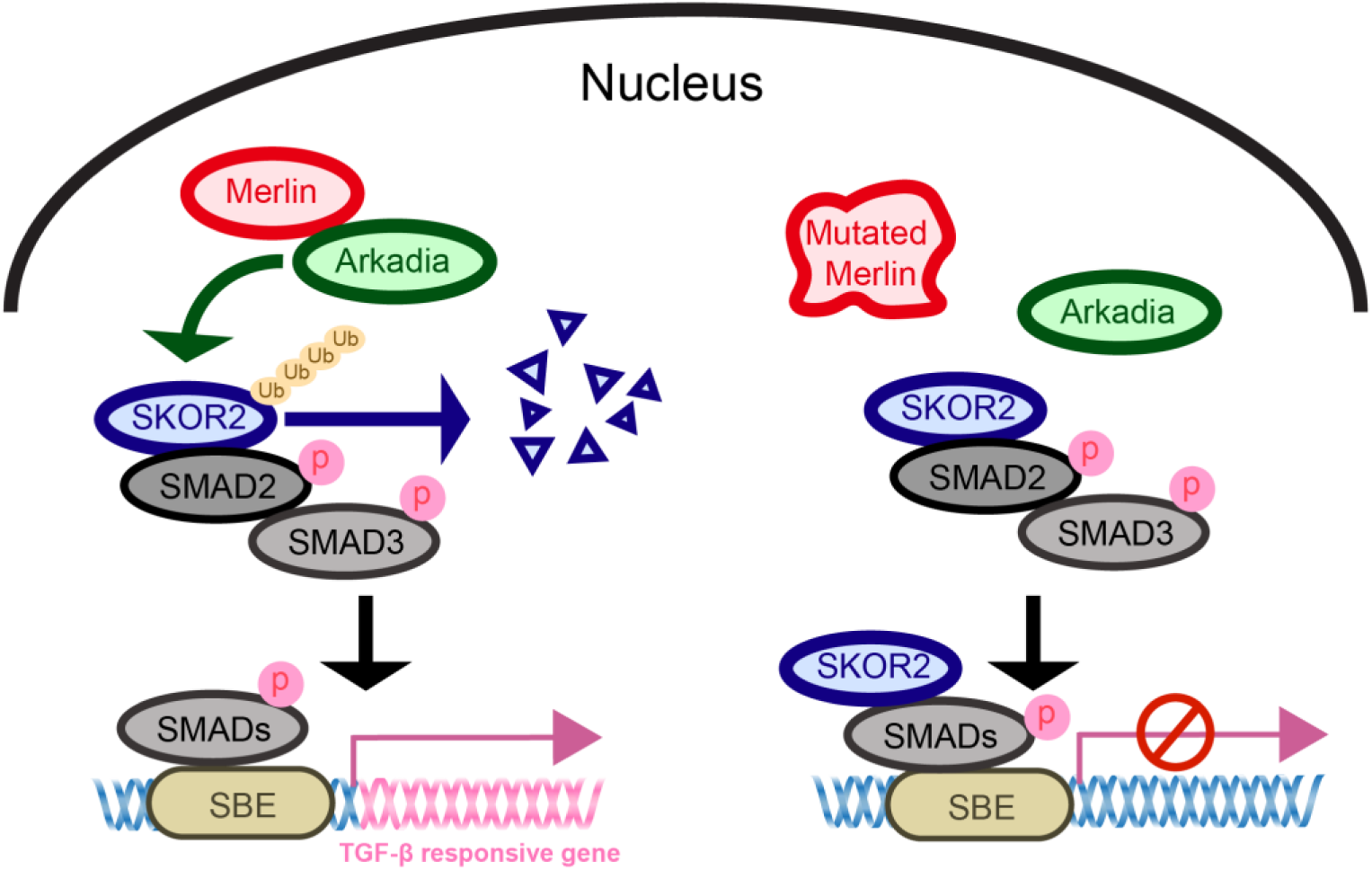
The model of interaction among merlin, Arkadia, and SKOR2 in responding to the SMAD-dependent TGF-β signaling pathway.

## Discussion

*NF2*-related schwannomatosis is a rare disorder caused by inherited or *de novo* mutations in the *NF2* gene, which lead to defects in the merlin protein(EVANS et al., 1992). The principal hallmark of *NF2*-related schwannomatosis is bilateral VS. Recent studies have elucidated many aspects of merlin function and suggest that it coordinates growth factor receptor signaling and cell adhesion. However, the molecular mechanisms underlying the effect of pathogenic *NF2* genetic variants remain unclear. This is due, at least in part, to the wide variety of signaling pathways. Merlin has been posited to control, including PI3K-AKT(Rong et al., 2004), RAC-PAK(Shaw et al., 2001), EGFR-RAS-ERK(Chiasson-MacKenzie et al., 2015; Curto et al., 2007), mTOR(James et al., 2009; López-Lago et al., 2009), and Hippo pathway(Hamaratoglu et al., 2006). Understanding the impact of different NF2 variants has been limited by the availability of model systems that faithfully recapitulate the human genetic landscape of VSs. Here in, we described a novel hiPSC-based SC model and showed that these SCs carrying NF2 patient specific variants could recapitulate the morphological and hyperproliferative phenotype seen *in vivo*. Combining this model with unbiased proteomic analysis, we were able to identify a novel interaction between merlin and the RING domain containing E3 ubiquitin ligase Arkadia and the SKI family transcriptional corepressor 2 (SKOR2).

SCs were successfully derived from three hiPSC lines – an isogenic pair of iPSC lines containing the NF2-associated p.L64P variant (NF2^L64P^) or the parental control line (NF2^WT^), as well as a NF2 patient iPSC line bearing a deletion in chromosome 22 that includes the *NF2* gene (NF2^+/−^) – following a predicted lineage transition. We firstly observed SOX10+ SCPs, which subsequently give rise to S100β+ SCs. Induction efficiency for SCs achieved ≥80%. These hiPSCs-derived SCs recapitulated phenotypes of *NF2* mutations. Specifically, SCs derived from NF2^L64P^ exhibited abnormal cell morphology compared to the isogenic parental control line (NF2^WT^). The morphology observed in the NF2^L64P^ SCs were reminiscent of those previously described(Gutmann et al., 1999). The NF2^L64P^ mutation is located in exon 2 and falls within the peptide region of merlin that binds directly to the molecular adaptor, paxillin(Fernandez-Valle C, 2002), which is involved in the recruitment of tyrosine kinases to focal adhesions, interactions with extracellular matrix, and actin organization(Schaller, 2001; Turner, 2000). This interaction with paxillin is important for establishing merlin localization and the regulation of cell morphology through the organization of actin(Fernandez-Valle C, 2002),(Brault E, 2001; Xu HM, 1998). Overall, our results supported that the NF2^L64P^ mutation result in aberrant cytoskeletal phenotypes.

Beyond its role in the cytoskeleton organization, merlin has been shown to interact with a variety of proteins. To examine how the NF2^L64P^ variant alters the merlin interactome, proteomic analysis using Co-IP followed by mass spectrometry was performed. Several proteins were identified that differentially bound (altered in the presence or quantity) to wild-type *NF2* and the p.L64P variant-bearing *NF2* in the hiPSCs-derived SCs. This dataset revealed many new protein candidates that interact with merlin in human SCs. We focused our analysis on two proteins, SKOR2 and Arkadia. Arkadia is an E3 ubiquitin protein ligase. Previously, merlin was reported to interact with another E3 ubiquitin protein ligase, CRL4 (DCAF1) (Li et al., 2010). While wild-type merlin interacts with Arkadia in hiPSCs-derived SCs, this interaction was disrupted by the missense mutation p.L64P. Arkadia was reported to ubiquitinate SKI family proteins in the SMAD-dependent pathway during TGFβ activation(Laigle et al., 2021; Sharma et al., 2011; Xu et al., 2021). Since SKOR2 was also identified in our proteomic analysis, upon further analysis, we found that SKOR2 accumulated in the nuclear fraction of NF2^L64P^ iPSC-derived SCs consistent with impaired turnover of SKOR2 by the UPS. In addition, we found that there was reduced levels of p-SMAD2/3 found in the NF2^L64P^ iPSC-derived SC nuclear lysates further supporting an impairment in TGFb signaling due to improper turnover of the transcriptional co-repressor SKOR2. This was validated using the SBE reporter assay which measures the activity of TGFβ/SMAD signaling pathway. Our results showed that the interaction between Merlin and Arkadia is associated with SKOR2 degradation, which enhances the response to TGFβ activation.

We next examined the effect of a deletion in chromosome 22 in which one copy of the NF2 gene is lost (NF2^+/−^) on SKOR2 levels and responses to TGFb signaling. Similar to what was seen for the NF2^L64T^ missense variant bearing iPSC-derived SCs, the NF2^+/−^ SCs had elevated SKOR2 levels in the nuclear protein fraction and reduced activity in response to the TGFβ activation – reduced p-SMAD2/3 levels and SBE activity – compared to NF2^WT^ SCs. In addition, the NF2^+/−^ SCs had elevated cellular proliferation levels as was seen with the NF2^L64P^ SCs. Interestingly, TGFβ signaling had been previously shown to be regulated by merlin. However, the effect of merlin was mediated through interactions with different components, e.g., PAK1 and 2(Wilkes et al., 2009), of the pathway in various cell types. Canonically, TGFb signaling leads to phosphorylation and activation of SMAD2/3 which, along with the SMAD4, interact with co-activators or co-repressors (e.g., SMAD7) to either activate or repress target gene transcription, respectively. Mota et al (2018) showed that loss of Merlin expression in breast cancer tissues was concordant with decreased SMAD7 expression leading to dysregulate TGF-β signaling pathway(Mota et al., 2018). Further, Cho et al. (2018) showed that Merlin activates non-canonical TGF-β type II receptor (TGFIIR) signaling leading to reduced TGF-β type I receptor (TGFIR) activity and abrogate its non-canonical oncogenic activity in mesothelioma(Cho et al., 2018). Thus, it appears that Merlin can target different portions of the TGFb signaling pathway to exert its tumor suppressor activity, including modulating SKOR2 stability, in different tumor types. Collectively, our findings proposed that Merlin functions as a tumor suppressor in hiPSC-derived SCs via interactions with Arkadia and SKOR2 to modulate the SMAD-dependent pathway in TGFβ signaling. This dysregulation of TGFβ signaling in the NF2^L64P^ and NF2^+/−^ iPSC-derived SCs could be responsible for driving the elevated cellular proliferation seen in these cells and, potentially, that seen during VS development.

## STAR Methods

### Cell culture and Schwann cell (SC) differentiation

Human induced pluripotent stem cells (hiPSCs) were maintained on the vitronectin-coated plate in the StemFlex medium (ThermoFisher). Media were changed daily.

Differentiation of hiPSCs toward SC followed a previous published protocol^17^ with modifications (Figure 1A). Briefly, hiPSCs were incubated in the NDM containing 1X N2, 1X B27, 0.005%BSA (Sigma), 2mM GlutaMAX (ThermoFisher), 0.11mM β mercaptoethanol (ThermoFisher), 3mM Chir99021(Reprocell), and 20 mM SB431542 (Reprocell) in advanced DMEM/F12 and Neurobasal medium (1:1mix) for 6 days prior to the incubation in NDM supplemented with 100 ng/ml NRG1 (Peprotech) for the Schwann cell precursor (SCP) induction. SCPs could be expanded and cryopreserved for the future usage. To further differentiate SCPs to SCs, SCPs were first incubated in SCDMI containing 1%FBS, 200ng/ml NRG1, 4mM forskolin (Sigma), 100nM retinoic acid (RA; Sigma) and 10ng/mL PDGF-BB (ThermoFisher) in DMEM/low glucose medium for 3 days. On the day 4, medium was replaced by SCDMII containing same ingredients as SCDMI without forskolin and RA. Two days later, cells were matured in SCM containing 1%FBS and 200ng/ml NRG1 for desired time.

### Patient-Derived Leukocytes

Assent and informed consent were obtained from a 12-year-old female with bilateral VS and her legal authorized representative, respectively, to collect and bank blood for research purposes, using a University of Miami Institutional Review Board-approved protocol (#20150637). The subject has a clinical diagnosis of NF2 and germline deletion of chromosome 22 that includes the *NF2* gene.

### Immunohistochemistry

Specimens were fixed in 4% paraformaldehyde for 30 min at RT with gentle shaking followed by three washes with PBS, 10 min each time. Blocking procedure used 10% desired serum in PBS with 0.1% triton X-100 for 30 min at RT. Subsequently, specimens were incubated with primary antibodies (Table S1) diluted in PBS with 3% goat or horse serum and 0.1% triton X-100. After washing with PBS for three times, specimens then were incubated with secondary antibodies at RT for one hr prior to three more washes with PBS. Finally, specimens were mounted using ProLong™ Gold Anti-fade mountant with DAPI (ThermoFisher). Images were taken using Keyence BZ-X series All-in-One Fluorescence Microscope.

### BrdU assay

Cell proliferation was measured using BrdU Cell Proliferation Elisa Kit (Abcam) following the manufacturer instruction. SCs were seeded on day 13 of the maturation stage with the same cell number. Cells were incubated with BrdU for 24 hrs prior to the measurement using a microplate reader.

### Cell surface measurement

Cells in 24 well plates were stained with phalloidin (Invitrogen) and mounted with ProLong™ Gold Anti-Fad mountant with DAPI. Images were taken using Keyence BZ-X series Fluorescence Microscope. Cell surface measurement and morphological analysis were taken in ImageJ. At least 50 cells were measured for each cell line.

### Cell treatment

SCs were induced with 2ng/ml TGFβ1 (PeproTech) before the SBE assay or protein isolation for the cytoplasmic and nucleus fractions.

#### Protein isolation and co-immunoprecipitation (Co-IP)

Total protein lysate was isolated using RIPA buffer (ThermoFisher) supplementary with protease inhibitors. Cytoplamic and nucleus proteins were isolated using NE-PER Nuclear and Cytoplasmic Extraction Kit (ThermoFisher). Protein lysates used for Co-IP were isolated using IP lysis buffer (ThermoFisher). All protein samples were quantified using Pierce BCA assay kit (ThermoFisher).

Co-IP was performed following the manufacture instruction of EZview™ Red Protein A Affinity Gel (Sigma-Aldrich) or Dynabeads™ Protein G Immunoprecipitation Kit (ThermoFisher). Briefly, antibody and protein lysate were incubated together for at least 1hr at 4°C to allow antibody-antigen complexes to form. Antibody-antigen complexes mix was then mixed with pre-washed gel beads at 4°C overnight. After three washes with lysis buffer, antibody-antigen complexes were eluted in SDS-PAGE sample buffer (Bio-Rad) for following applications, e.g., SDS-PAGE analysis and Western blots.

#### TMT labeling and mass spectrometry

After Co-IP, proteins were eluted in 0.2M glycine (pH 2.5) and dried in the vacuum concentrator. The six samples (n=3 each for the WT & mutant, with each replicate from the same genotype representing a unique culture) were prepared for TMT labeling and mass spectrometry (MS) with the EasyPrep™ MS sample prep kit (ThermoFisher). Subsequently, the digested peptides were incubated with TMT labels 131C, 132N, 132C, 133N, 133C, and 134N, quenched with hydroxylamine followed by the peptide purification. Purified labeled peptides were dried to completion and resuspended in 10 µl of 2% acetonitrile with 0.1% formic acid. Peptide identification from MS was completed by the Ophthalmology mass spectrometry core facility in the University of Miami Miller School of Medicine.

#### Bioinformatics

RAW files from the MS were imported into Proteome Discoverer (ver. 3.0; TFS) and analyzed using the default TMT workflow (minus the first 10 labels of the 16-plex kit, which were used for another experiment). As the first step in this workflow, a conceptually translated human genome (as a fasta file; give details about the genome) was queried using the default conditions from Proteome Discoverer’s Sequest-derived algorithm. Both quantitative (TMT-labeled peptides) and semi-quantitative (presence-absence) data were exported as .csv files and imported into JMP® Pro (ver. 17; Cary, NC, USA). All proteins were scaled by Proteome Discoverer to where the mean of the six samples was 100; this ensured that high abundance proteins did not bias the multivariate analyses outlined below.

However, this scaling step does not ensure that each sample yields comparable data. To demonstrate this, the overall mean TMT signal was assessed across all six samples, and it was found to differ significantly (*p*<0.01) among them; some samples consistently yielded higher protein concentrations than others, despite having labeled the same amount of protein. To correct for this, the concentrations of the individual proteins were normalized to the global mean of the respective sample. For instance, if sample A yielded an overall mean TMT level of 150 across all 262 labeled peptides and sample B yielded 75 across these same peptides, the individual peptide concentrations of samples A and B would be divided by 150 and 75, respectively, to ensure that laboratory benchwork-associated bias did not influence results. Upon undertaking this normalization step, the mean protein level was reduced from the Proteome Discoverer default of 100 to 1.

As the simplest means of uncovering treatment-responsive proteins, proteins found in all three replicates of one treatment and in no samples of the other were uncovered (i.e., both WT-only & mutant-only). As a more common means of identifying differentially concentrated proteins (DCPs), JMP’s response screen was used. This analysis features an FDR-adjustment of the alpha level to where the chance of making a false-positive statistical error on account of having made some many comparisons is reduced. Only proteins that were both significantly differentially concentrated at an FDR-adjusted alpha of 0.05 and that differed by at least 2-fold between treatments were considered to represent DCPs.

As a more global means of characterizing treatment effects on the partial IP-proteome, both principal components analysis (PCA) and multi-dimensional scaling (MDS) were performed with the subset of 262 TMT-labeled peptides. To determine whether there was a multivariate difference between the proteomes of samples of the two treatments, a non-parametric multivariate ANOVA (NP-MANOVA) was undertaken using the coordinates from the first three MDS dimensions (stress=0.08) as the model Y’s. This analysis was used because standard MANOVA cannot be undertaken with wide datasets (i.e., more analytes than samples), and an alpha of 0.05 was set *a priori*. Partial least squares was used simultaneously to generate a model such that the misclassification rate could be calculated (a secondary means of gauging the degree of difference between the partial proteomes of the two treatments).

### Western blots

SDS-PAGE was performed using the Bio-Rad Mini-PROTEAN Tetra system. Western blots were performed following standard procedures. Antibodies used in this study were listed in the table S1. Secondary antibodies conjugated with horseradish peroxidase (HRP) were used. Development of images used SuperSignal™ West Femto Maximum Sensitive Substrate (ThermoFisher). Images were taken using ChemiDoc Imaging System (Bio-Rad).

### SBE assays

SBE assay was performed using SBE reporter kit (BPS Bioscience) following the product general protocol. Cells were transfected using Lipofectamine 2000 (Invitrogen) and treated with TGFβ1 for 8 hours. Two-Step Luciferase (Firefly and Renilla) assay system (BPS Bioscience) was used to measure the SBE reporter activity. Firefly luciferase readouts were normalized with Renilla readouts prior to the statistical analysis.

### Statistical analyses

T-test and One-way ANOVA were performed using Prism 10. *p-*value <0.05 was deemed to be significant. Sample size was ≥3 in every experiment, measurement, and statistical analysis.

## Acknowledgements

We acknowledged the service provided by the Ophthalmology mass spectrometry core facility in the University of Miami Miller School of Medicine. The work was supported by NIH grants of R01DC017264, R01DC005575, R01DC012115, and NIDCD R25 DC020726 to XL, R21DC019450 to P-CT, and K08DC017508 and Sylvester K-supplement to CTD.

## Author contribution

P-C. T.: conceptualization, performing experiments, data analyses, and wrote the manuscript. S.U.: performing experiments. A.B.M.: performing experiments, data analyses, and editing the manuscript. C.D.C.: performing experiments. O.R.B.: performing experiments. C.D.: patient recruitment, patient screening, providing materials and editing the manuscript. D.M.D.: conceptualization and editing the manuscript. X.L.: providing resources. All authors reviewed and approved the final draft of manuscript.

