## Supplemental information for "Interactions among Merlin, Arkadia, and SKOR2 mediate NF2-associated Schwann cell proliferation in human"

### Supplementary information

A

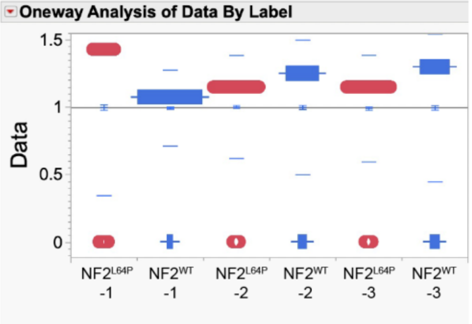


**Means and Std Deviations**

| Level | Number | Mean | Std Dev | Std Err Mean |
|---|---|---|---|---|
| NF2$^{L64P}$-1 | 1076 | 0.9997345 | 0.655074 | 0.0199703 |
| NF2$^{WT}$-1 | 1076 | 0.9953232 | 0.2822158 | 0.0086035 |
| NF2$^{L64P}$-2 | 1076 | 1.0041448 | 0.3821235 | 0.0116492 |
| NF2$^{WT}$-2 | 1076 | 1.0002323 | 0.5000581 | 0.0152445 |
| NF2$^{L64P}$-3 | 1076 | 0.9913259 | 0.396073 | 0.0120745 |
| NF2$^{WT}$-3 | 1076 | 0.9969584 | 0.5487303 | 0.0167283 |

B

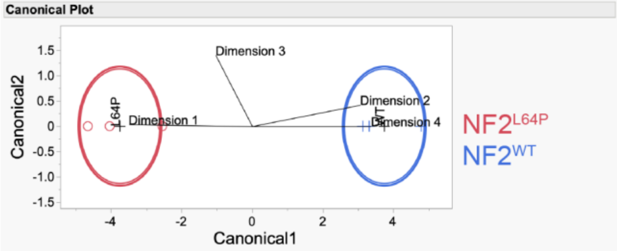


**Canonical Details**

Canonical Details calculated from the overall pooled within-group covariance matrix.

| Eigenvalue | Percent | Cum Percent | Canonical Corr | Likelihood Ratio | Approx. F | NumDF | DenDF | Prob>F |
|---|---|---|---|---|---|---|---|---|
| 21.0803958 | 100.0000 | 100.0000 | 0.97709312 | 0.04528904 | 5.2701 | 4 | 1 | 0.3144 |

| Test | Value | Exact F | NumDF | DenDF | Prob>F |
|---|---|---|---|---|---|
| Wilks' Lambda | 0.045289 | 5.2701 | 4 | 1 | 0.3144 |

C

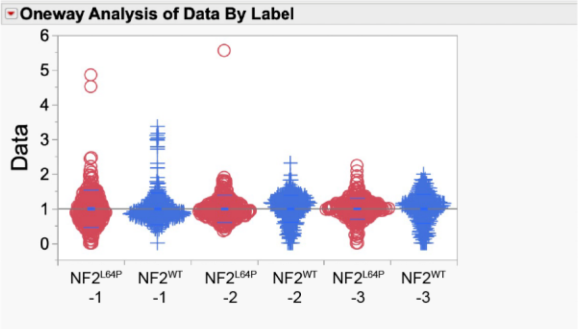


**Means and Std Deviations**

| Level | Number | Mean | Std Dev | Std Err Mean | Lower 95% |
|---|---|---|---|---|---|
| NF2$^{L64P}$-1 | 263 | 1.0007996 | 0.536709 | 0.0330949 | 0.9356338 |
| NF2$^{WT}$-1 | 263 | 0.9979366 | 0.4600378 | 0.0283671 | 0.94208 |
| NF2$^{L64P}$-2 | 263 | 0.9968561 | 0.3947846 | 0.0243435 | 0.9489223 |
| NF2$^{WT}$-2 | 263 | 0.9984316 | 0.3895008 | 0.0240176 | 0.9511394 |
| NF2$^{L64P}$-3 | 263 | 0.9981664 | 0.2989212 | 0.0184323 | 0.9618721 |
| NF2$^{WT}$-3 | 263 | 0.9991644 | 0.4163762 | 0.0256749 | 0.9486091 |

Figure S1. Mass spectrometry data analyses. A. Normalization of total 1076 proteins with presence-absence data. B. Presence-absence protein data separate samples between NF2$^{WT}$ and NF2$^{L64P}$ in the non-parametric multivariate analysis of variance (MANOVA). C. Normalization of total 263 proteins with TMT-labeling.

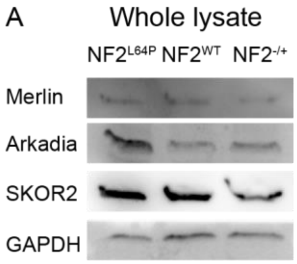

Figure S2. Western blots of whole lysates isolated from hiPSC-derived SCs.

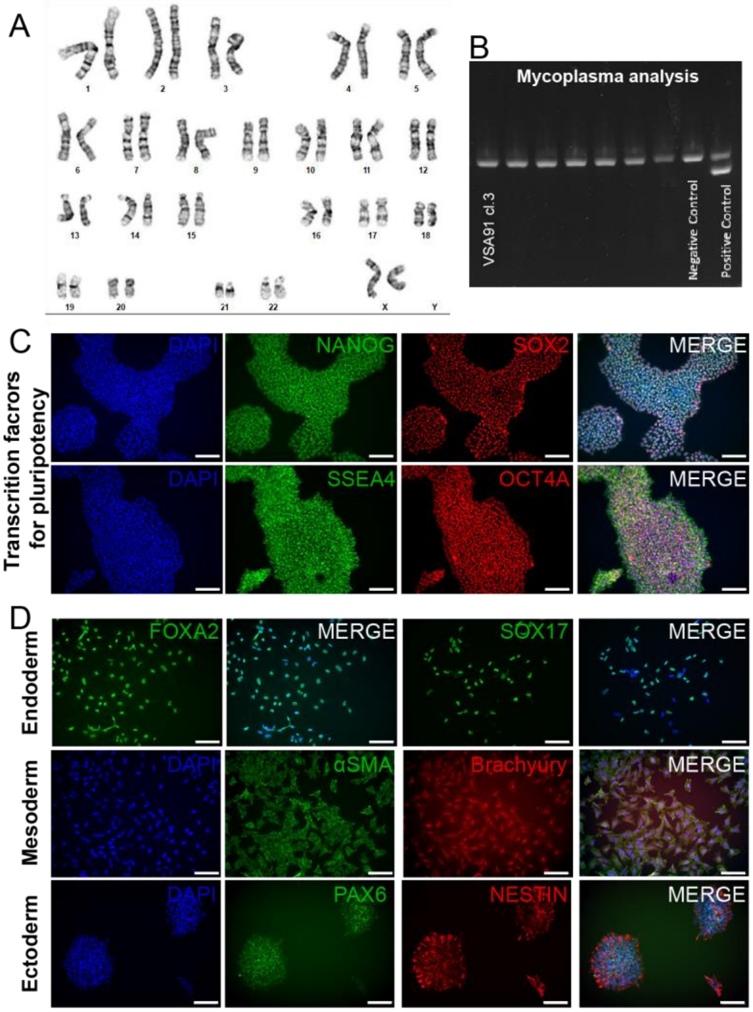


Figure S3. Characterization of a patient-derived $NF2^{+/-}$ hiPSC line, VSA91. A. G-band karyotype showed normal chromosome structure. B. Agarose gel image indicated the cell line was negative from mycoplasma. C. Representative immunohistochemistry (IHC) images of markers for pluripotency. D. Representative IHC images for the trilineage differentiation.

Table S1. Antibodies list

| Antibody | Isotype | Dilution | Application | Manufacturer |
|---|---|---|---|---|
| SOX10 | Rabbit IgG | 1:50 | IHC | Abcam |
| S100β | Rabbit IgG | 1:100 | IHC | Abcam |
| NeuN | Rabbit IgG | 1:500 | IHC | Abcam |
| NANOG | Rabbit IgG | 1:100 | IHC | Invivogen |
| SOX2 | Mouse $IgG_1$ | 1:50 | IHC | BD Bioscience |
| SSEA4 | Mouse $IgG_3$ | 1:100 | IHC | StemCell Technology |
| OCT4A | Rabbit IgG | 1:200 | IHC | Cell Signaling Technology |
| FOXA2 | Mouse $IgG_{2a}$ | 1:200 | IHC | Abcam |
| SOX17 | Goat IgG | 1:100 | IHC | R&D Systems |
| αSMA | Mouse $IgG_{2a}$ | 1:200 | IHC | Abcam |
| Brachyury | Rabbit IgG | 1:200 | IHC | Abcam |
| PAX6 | Rabbit IgG | 1:100 | IHC | Abcam |
| Nestin | Mouse $IgG_1$ | 25 μg/ml | IHC | R&D Systems |
| Merlin | Rabbit IgG | 1:1000 | WB | Cell Signaling Technology |
| Merlin | Mouse $IgG_{2b}$ | 1:200 / 5 μg | WB/IP | Santa Cruz |
| Arkadia/RNF111 | Rabbit IgG | 1:1000 / 5 μg | WB/IP | Invitrogen |
| SKOR2 | Rabbit IgG | 1:1000 | WB | LS Bio |
| Ubiquitin | Mouse $IgG_1$ | 1:100 | WB | Invitrogen |
| p-SMAD2/3 | Rabbit IgG | 1:1000 | WB | Cell Signaling Technology |
| GAPDH | Mouse $IgG_1$ | 1:100 | WB | Santa Cruz |
| TBP | Rabbit IgG | 1:1000 | WB | Cell Signaling Technology |
